# Nitrite accumulation and the associated anammox bacteria niche partitioning in marine sediments

**DOI:** 10.1101/2022.08.18.504054

**Authors:** Rui Zhao, Andrew R. Babbin, Desiree L. Roerdink, Ingunn H. Thorseth, Steffen L. Jørgensen

## Abstract

By consuming ammonium and nitrite, anammox bacteria form an important functional guild in nitrogen cycling in many environments including marine sediments. Recent studies have shown that anammox bacteria can consume most of the upwardly diffusing ammonium from deep anoxic sediments; however, their impact on the other important substrate nitrite has not been well characterized. Here we show niche partitioning of two anammox families emerges in a 2.4-m long mostly anoxic sediment core retrieved from the Nordic Seas. We document high abundances (~10^6^ cells g^−1^) of anammox bacteria in most investigated sediment layers, with two distinct anammox abundance maxima in two nitrite consumption zones. Between the two anammox abundance maxima, nitrite accumulates as observed in other marine sediment sites and aquatic environments, indicating anammox bacteria play a fundamental role in modulating the nitrite distribution. Anammox bacteria in the upper nitrite consumption zone are dominated by the *Candidatus* Bathyanammoxibiaceae family, while *Ca*. Scalinduaceae family dominate at the lower zone. A high-quality representative *Ca*. Bathyanammoxibiaceae genome is recovered, which, comparing to *Ca*. Scalindua sediminis, the representative of Scalinduaceae in marine sediments, has fewer high-affinity ammonium transporters and lacks the capacity to access alternative substrates or energy sources such as urea and cyanate. These features may restrict *Ca*. Bathyanammoxibiaceae to conditions of higher ammonium concentrations or fluxes, and therefore drive the observed niche partitioning. These findings improve our understanding about nitrogen cycling in marine sediments by revealing the association between nitrite accumulation and niche partitioning of anammox bacteria.

## Introduction

The cycling of nitrogen in ecosystems is intricately controlled by a network of processes mediated by microbes. Dinitrogen gas (N_2_) is the most abundant nitrogen species but not available to most organisms, while most of the other nitrogen species in the nitrogen cycle are bioavailable but much scarcer. In an ecosystem, new bioavailable (or fixed) nitrogen is generated by diazotrophy, and can be converted back to N_2_ by two nitrogen loss processes: denitrification and anaerobic ammonium oxidation (anammox) [see reviews in e.g. [1,2]]. The latter two anaerobic metabolisms are generally favored in low-oxygen environments, either the ocean’s pelagic oxygen minimum zones or benthic sediments [2–4]. Although whether the fixed nitrogen sources and sinks are balanced [5,6] or unbalanced [7] is still debated, it is broadly accepted that the benthos represent quantitatively the most important fixed nitrogen loss system in the marine environment [6,8,9]. As a consequence, the benthic environment plays a crucial role in regulating the abundance of bioavailable nitrogen across the marine realm. Nitrite is a crucial substrate or intermediate of both nitrogen loss processes [10,11], the availability of which exerts a profound control on the magnitude of nitrogen loss [12]. However, nitrite rarely accumulates to as high a level as nitrate and ammonium in marine sediments, leading the presence and cryptic cycling of nitrite in this vast environment to be largely overlooked.

Since its discovery in the marine environment two decades ago [13], anammox has been shown to dominate fixed nitrogen loss [e.g., [14–16]] and therefore control global ocean productivity [17]. Among the previously recognized marine anammox bacteria (affiliated with the families *Candidatus* Brocadiaceae and *Candidatus* Scalinduaceae), members of Scalinduaceae have been consistently detected in marine sediments [14,16,18–20], with several enrichment cultures obtained from coastal sediments [e.g., *Ca*. Scalindua japonica [21] and *Ca*. Scalindua profunda [22]]. Recently, a new family of anammox bacteria (i.e., *Candidatus* Bathyanammoxibiaceae) has been discovered in marine sediments, which in some cores can significantly outnumber their counterparts of Scalinduaceae [23]. How *Ca*. Bathyanammoxibiaceae interacts with *Ca*. Scalinduaceae in marine sediments and what influence they have on nitrite transformations are still unclear.

To improve our understanding about anammox bacteria and their biogeochemical roles in marine sediments, we combine biogeochemical, microbiological and genomic approaches to study the relationships between the distribution of dissolved nitrogen species and anammox bacteria. We first identify a widespread phenomenon of nitrite accumulation in the nitrate-depletion zone in diverse marine sediment systems: the continental slope, mid-ocean ridges, and also hadal sediments. Through high-resolution analyses of microbial communities in a sediment core from the Arctic Mid-Ocean Ridge (AMOR), we observe niche partitioning of anammox bacteria between the families *Ca*. Scalinduaceae and *Ca*. Bathyanammoxibiaceae that are prevalent in marine sediments. Based on the newly generated high quality anammox genomes, we also propose the likely underlying genetic mechanisms for the niche partitioning and thus for the accumulation of nitrite.

## Results and Discussion

### General geochemical context of GS14-GC04

In this study, we focused on a 2.4-m-long core (GS14-GC04, called GC04 henceforth) retrieved from a 1050-m deep seamount 50 km west of the Jan Mayen hydrothermal vent fields (Fig. 1A) where white smoker hydrothermal vents were reported [24,25]. The retrieved sediments of GC04 were oligotrophic, with measured total organic content lower than 0.5 wt% (Fig. S2A). Organic nitrogen content was measured to be in the range of 0.06-0.11% (Fig. S1A) and thus the calculated carbon to nitrogen ratio (C/N) fell generally in the range of 2 – 4 (Fig. S1B). Oxygen was measured to be only 15 μM at the top of the recovered core and was depleted within 23 cm below the seafloor (cm bsf). Below the depletion depth of oxygen, dissolved Mn was measured to accumulate in the porewater (Fig. 2B). Porewater pH was measured between 7.6 and 7.8 (Fig. S1C), similar to those in other AMOR cores [18]. Dissolved Fe was not detected (Fig. S1D) and sulfate was measured to be constant and at bulk seawater values (with uncertainty <0.5%) throughout the core (Fig. S1E), indicating that the reduction of Fe and sulfate are not important redox reactions in the recovered sediments. GC04 exhibited higher concentrations of dissolved inorganic carbon (DIC) and dissolved Mn (Fig. S2) than the four AMOR cores without significant hydrothermal influences [18], indicating higher organic matter degradation activity in GC04 probably caused by higher seafloor organic matter load due to its shallower water depth than other cores. Despite the uppermost sediments potentially been lost during coring (see Supplementary Note 1), the oxygen penetration depth was shallower than the non-hydrothermal sites and does not affect our interpretation of deeper anaerobic microbes and their metabolisms.

**Fig. 1.**
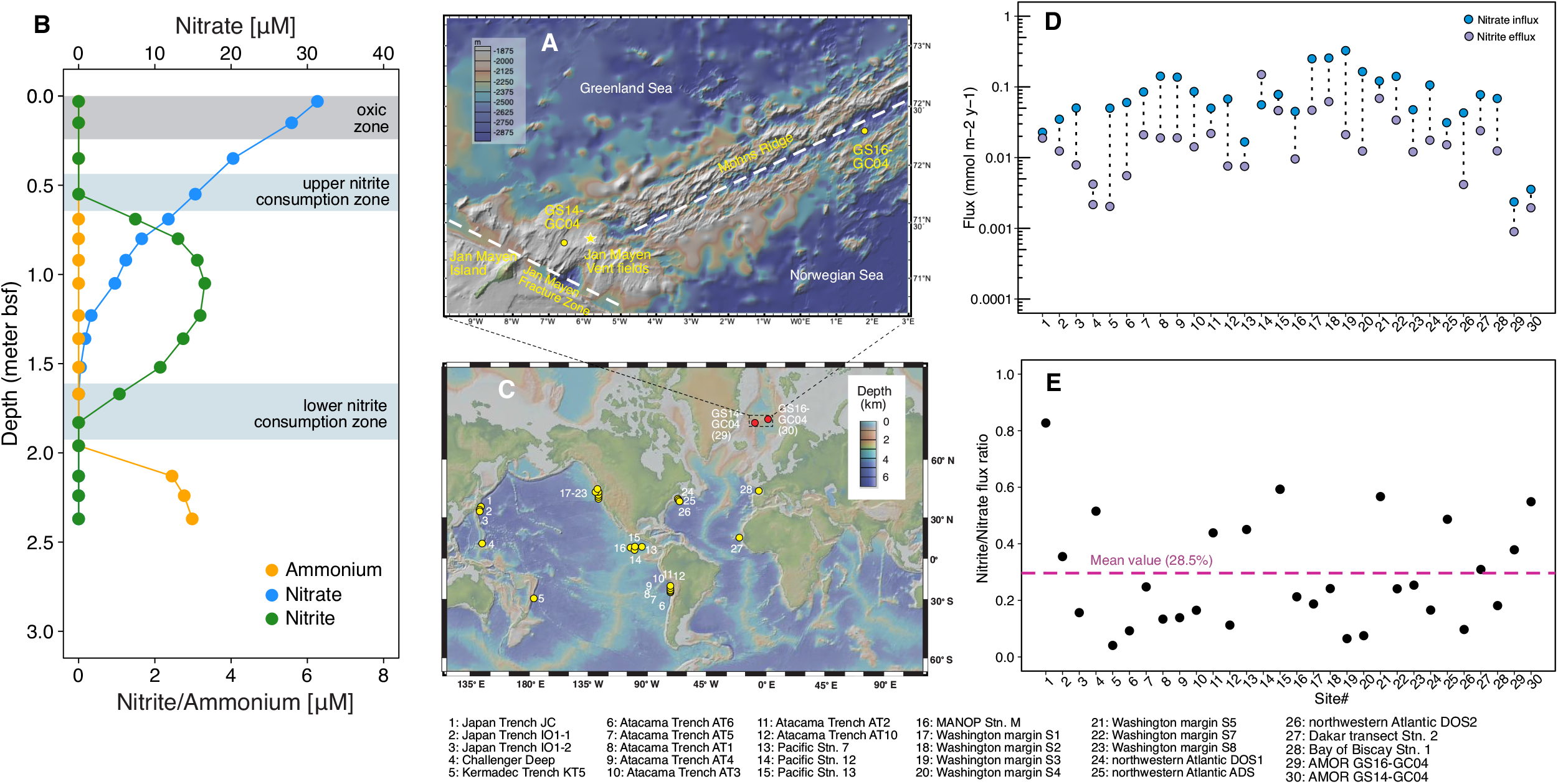
The occurrence of nitrite accumulation in sediment porewater of the Arctic Mid-Ocean Ridge and other locations. **(A)** Bathymetric map showing two coring site (GS14-GC04 investigated in this study and GS16-GC04 investigated in Ref. 18) in the Arctic Mid-Ocean Ridge area where nitrite accumulation was observed. Also highlighted are the Jan Mayen Fracture Zone and the Mohns Ridge, as well as the Jan Mayen hydrothermal vent field (red star). The insert map showing the study location in the broad Arctic area. **(B)** Porewater profiles of nitrate, nitrite and ammonium in GC04. For visibility, 5 times the measured nitrite concentrations are shown. The oxic zone and two net nitrite consumption zones are highlighted by horizontal bands. (**C**) Sediment locations where the accumulation of nitrite in the porewater was detected (See Fig. S3 for the porewater nitrite and nitrate profiles of individual sites). The two AMOR sites are shown in red circles, while other sites are shown in yellow circles. Maps in (**A**) and (**C**) were generated with GeoMapApp v. 3.6.14 (www.geomapapp.org), using the default Global Multi-Resolution Topography Synthesis basemap. (**D**) Nitrate influx and (combined upward and downward) nitrite efflux in the nitrate-depletion zone. The paired fluxes for each site are connected with a black dotted line. (**E**) Calculated nitrite/nitrate flux ratio for the individual sites. The horizontal line denotes the mean value of the 30 sites.

**Fig. 2.**
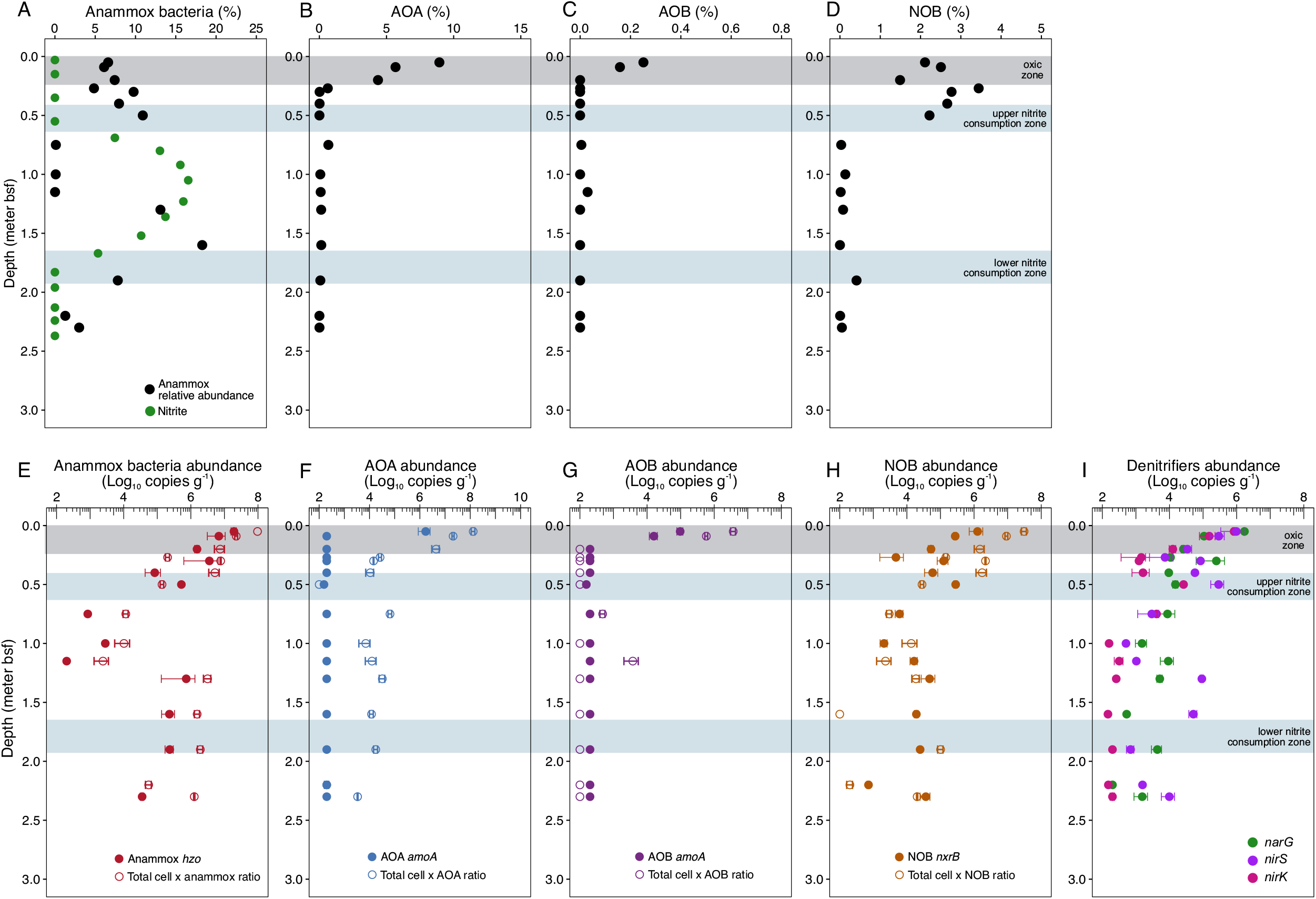
Abundances of anammox bacteria (A), ammonia-oxidizing archaea (AOA, B), ammonia-oxidizing bacteria (C), nitrite-oxidizing bacteria (NOB, D), and denitrifying bacteria (E) in GC04. Filled circles indicate the absolute abundances of these groups determined by qPCR using specific primers targeting their diagnostic genes, while open circles in (A, B, and C) denote the absolute abundances of anammox bacteria, AOA, and AOB calculated as the product of the total cell numbers (shown in Fig. S4A) and their respective relative abundance in the total community.

### Accumulation of nitrite in the nitrate-depletion zone

This core features an extended nitrogenous zone. Nitrate was found to decrease with depth and was depleted around 130 cm bsf (Fig. 1B). Ammonium was not detected in the porewater until 213 cm bsf, well below the nitrite and nitrate depletion depths (Fig. 1B). Therefore, this core has an extended nitrate-ammonium transition zone (NATZ), spanning a full meter (i.e., 120 – 220 cm bsf).

Unlike most sediment cores where nitrite is generally undetectable throughout all measured depths, nitrite in GC04 accumulated around the nitrate-depletion zone (70 – 160 cm bsf), with a concentration maximum (~ 3 μM) at 105 cm bsf. A similar nitrite profile, albeit of lower magnitude, was also detected in the nitrate-depletion zone of GS16-GC04 [26] among the four AMOR cores reported in [18]. By searching published literature, we found that such accumulation of nitrite around the nitrate-depletion zone was also detected in 28 additional globally distributed sediment cores (Fig. 1C; See the detailed nitrite and nitrate profiles in individual cores in Supplementary Figure S3). Such accumulation was mainly detected in sediments on the continental slopes [[27–30]], along the mid-ocean ridges [31] of the Pacific and Atlantic Oceans, and within hadal trenches in the Pacific [14,32,33] (Fig. 1B). Notably, most of these sites also feature a nitrate-ammonium transition zone (NATZ), where the anammox reaction occurs [14,18], suggesting a potential link between anammox bacteria and the observed nitrite accumulation. The nitrite accumulation was likely not detected in the upper few meters of sediments of (i) continental margins because nitrate penetration is too shallow to be properly resolved without dedicated microscale measurements, or (ii) abyssal plains because high concentrations of nitrate and O_2_ are present in the porewater throughout the entire sediment columns [e.g., [33–36]]. Through this comparison, it is likely that the observed nitrite accumulation in sediments of continental slopes, mid-ocean ridges, and hadal trenches is tightly associated with low concentrations of nitrate within the extended nitrogenous zone, which in turn is caused by moderate levels of organic matter flux.

The nitrite accumulation in the lower nitrate zone was also detected in other stratified aquatic environments like the water columns of the Black Sea [15,37] and Golfo Dulce [38], the freshwater Lake Tanganyika [39], hypersaline Lakes Vanda and Bonney [40,41] in the McMurdo Dry Valleys in Antarctica, river and estuary sediments [42,43], subtropical mangrove sediments [44], and denitrifying biofilms in wastewater treatment plants [45]. The observations indicate that the accumulation of nitrite within the nitrogenous zone is likely a widespread phenomenon in aquatic environments that harbor redox gradients.

### Accumulated nitrite only accounts for a small fraction of the consumed nitrate

The observed nitrite maxima in marine sediments (up to 8 μM as reported in [30]) are comparable or higher than those measured in oxygen deficient zones [e.g., [46,47]], but are generally lower than the concomitant nitrate concentrations, indicating that nitrite is only a minor inorganic nitrogen species in the sediments. Yet, nitrite is a central metabolite for many microorganisms, and the low concentrations only imply its fast turnover is well coupled in the environment rather than it is an unimportant metabolite [48].

Nitrite accumulation in the nitrate-depletion zone also indicates that some of the detected nitrite can diffuse both upward and downward and support two distinct zones (e.g., above and below the nitrate depletion depth) that harbor intensified nitrite consumption. By calculating the nitrate influx and the total (the sum of the upward and downward) efflux of nitrite from the nitrate-depletion zone, we found that at all but one site (site #14, Pacific Station 12 reported in [30]), the nitrate flux is higher than that of the combined nitrite flux at all sites (Fig. 1D). Because of this, the calculated ratio of nitrate to nitrite flux at all but one site is less than 0.6 (Fig. 1E), with the average ratio of 0.285±0.196 (mean ± stdev, with the median value of 0.241), suggesting that nitrite flux only accounts for on the order of a quater of the nitrate flux consumed within the nitrate-depletion zone and that the majority of the nitrate diffused into that zone is lost by further reduction to unmeasured gaseous compounds (e.g., N_2_).

### The prevalence of anammox bacteria throughout the core

To elucidate whether microbes play a role in controlling the observed nitrite accumulation in marine sediments, we performed 16S rRNA gene amplicon sequencing for 13 sediment layers of GC04. We noted the prevalence of putative anammox bacteria (affiliated to both families *Ca*. Scalinduaceae and *Ca*. Bathyanammoxibiaceae [23]) in most layers. Anammox bacteria, notoriously slow growers [49], are sizable contributors of the communities, accounting for 6% of the total community in the uppermost sediments in the oxic zone, and increasing to a first peak of 11% of the total community in the upper nitrite consumption zone (Fig. 2A). After a major collapse in the interval of 75 – 120 cm bsf, the relative abundance of anammox increases again and reaches the second peak of a full ~18% of the total community within the second nitrite consumption zone before again decreasing in deeper sediments (Fig. 2A). The second peak is roughly within the broad NATZ.

To check whether the relative abundance changes of anammox bacteria are caused by growth/decay of other taxa, we tracked the absolute abundances of anammox bacteria using two complementary methods: (i) qPCR of the functional gene *hzo*, which encodes hydrazine dehydrogenase, the ultimate step of the anammox metabolism and therefore a diagnostic gene for anammox bacteria, and (ii) calculation as the product of the total cell abundance (estimated as the sum of the 16S rRNA genes presented in Fig. S4A) and the relative abundances given by the 16S rRNA gene amplicon sequencing. As shown for other AMOR cores [18], results of the two methods generally agree (Fig. 2E), indicating the major anammox clades are accounted for in this analysis. The prevalence of anammox bacteria in the upper and lower portions of GC04 is corroborated by their high absolute abundances in the range of 10^6^–10^8^ cells g^−1^ wet sediment, while relatively lower abundances of 10^2^–10^4^ cells g^−1^ are detected in the middle section of the core (75–120 cm bsf) (Fig. 3E). Contrasting with previous reports that sediment anammox bacteria are restricted within the NATZ [18,23], our results suggest that the distribution of anammox bacteria is more complex than previous studies would imply.

**Fig. 3.**
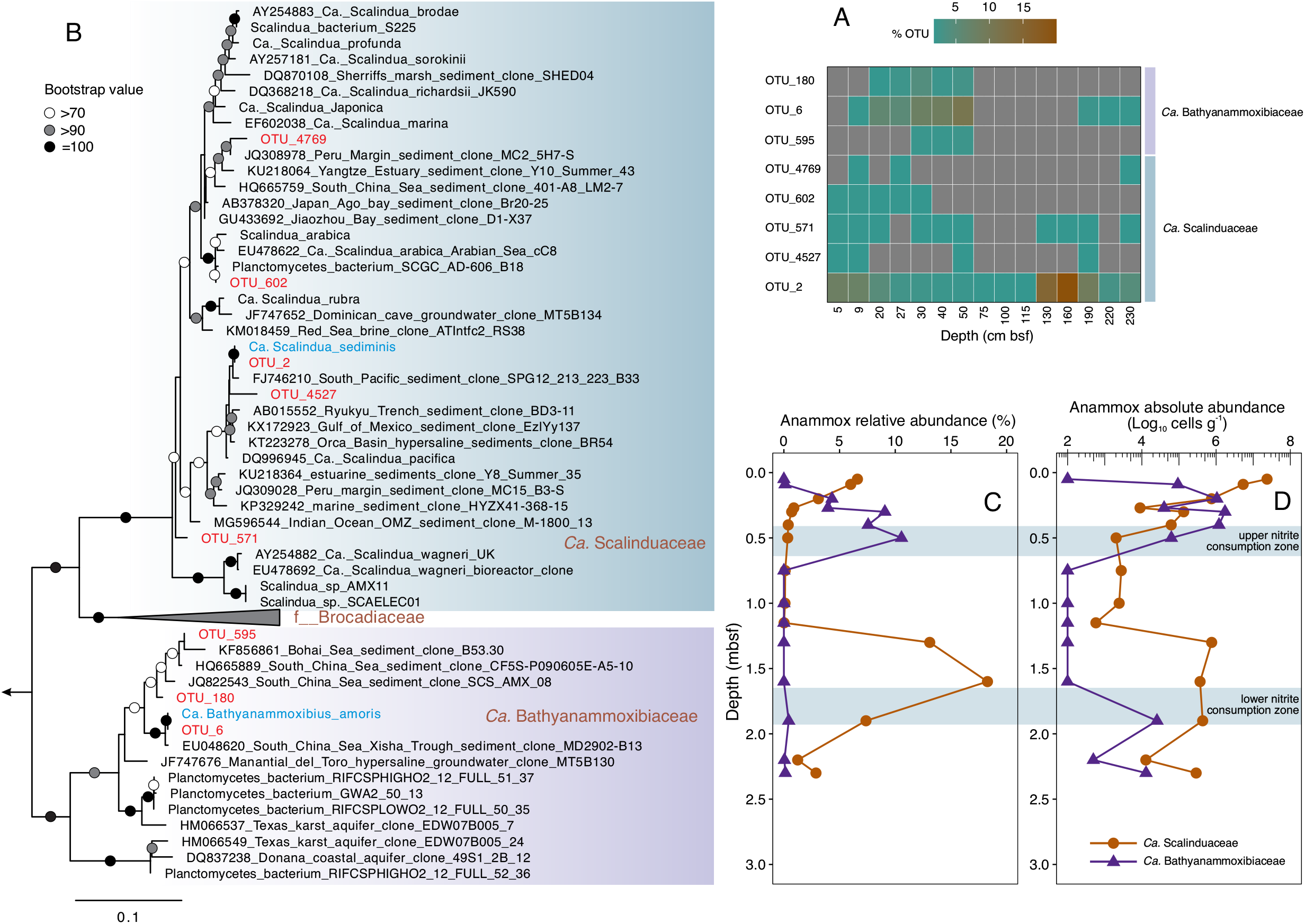
Distribution, phylogeny, and abundances of anammox bacteria in GC04. **(A)** Heatmap showing the occurrence of eight anammox OTUs in the investigated sediment layers in core GC04. **(B)** Maximum-likelihood phylogenetic tree of anammox bacteria. Anammox bacteria OTUs (97% identity cutoff) recovered from GC04 are highlighted in blue and red colors. The bar indicates estimated sequence divergence per residue. (**C**) Relative abundances of the anammox families (*Ca*. Scalinduaceae and *Ca*. Bathyanammoxibiaceae) throughout core GC04, as assessed by the 16S rRNA gene amplicon sequencing. (**D**) Absolute abundances of the two anammox families, calculated as the product of the total cell numbers times the relative abundances of them shown in (**C**).

The presence of anammox bacteria (mainly affiliated with *Ca*. Scalinduaceae) in the oxic zone (Fig. 2E, with up to 20 μM O_2_) of GC04 is interesting. Although early bioreactor studies have shown that 1 μM O_2_ reversibly inhibits the anammox metabolism [50], anammox bacteria and activity have been detected in seawater with up to 25 μM O_2_ [15,51], which may be facilitated by associating with particles [15,52] and the microenvironments therein [53]. Particles and colonized surfaces are widespread in marine sediments, which can harbor anoxic microniches to greatly expand the anoxic habitats even in bulk oxygenated environments [54,55]. Therefore, increased anoxic microenvironments in hydrothermal sediments, which typically have larger grain size than typical sediments [56], could enable the presence of anammox bacteria in the bulk oxic surface sediments. Alternatively, the anammox bacteria detected in the oxic zone could be dormant. Nevertheless, the detection of anammox bacteria in the surface sediments do confirm the hypothesis that anammox bacteria thriving in subsurface NATZs were seeded from surface sediments [18]. Future studies are required to assess anammox bacteria activity and contribution to cryptic nitrogen transformations in the shallow oxic sediments.

### Role of anammox bacteria for ammonium and nitrite consumptions in the interval between OATZ and NATZ (the nitrogenous zone)

Ammonium is the major nitrogen species present in sediment porewaters of continental shelves and slopes. In these sediments, ammonium is mainly produced from organic nitrogen degradation and can be consumed by biological activities such as aerobic ammonia oxidation and anammox and also biological re-assimilation, albeit the latter should be minimal due to the extremely slow microbial turnover rates. Previous studies have shown that AOA prevail in the oxic zone [26,35] and anammox bacteria in the NATZ [18], respectively, which may be the major ammonium consumers in their major niches. In GC04, despite continuously ammonium release from organic matter degradation as evident by the increasing DIC concentrations with depth (Fig. S2A), ammonium was not detected until both nitrate and nitrite were depleted from the porewater (Fig. 1B), suggesting active ammonium consumption throughout the upper 1.8 m sediments of GC04. However, ammonium consumption in the sediment interval between the OATZ and the NATZ [i.e., the nitrogenous zone [57]] is still unclear.

To better understand the relative importance of anammox bacteria for ammonium consumption in GC04, we examined, in addition to anammox bacteria themselves, the distribution (i.e., both the relative and absolute abundances) of aerobic ammonia-oxidizing archaea (AOA) and bacteria (AOB) in this core using the two microbial quantitative methods described above. Consistent with their requirement of oxygen [58,59], both AOA (affiliated with the class Nitrosopumilales of the phylum Thaumarchaeota [60,61]) and AOB are mainly detected in the upper 10 cm sediments of the oxic zone (Fig. 2B and 2C). While AOB (<0.3% of the total communities throughout the core) seem to be restricted to the oxic zone (Fig. 3G), AOA are detected not only in the oxic zone but also in deeper anoxic sediments (yet <0.7% of the total deep communities) by 16S rRNA gene amplicon sequencing, although AOA *amoA* gene-based qPCR only detects AOA in the oxic zone (Fig. 2F). The discrepancy may be attributed to the possibility that the qPCR primers of the AOA *amoA* gene assays may fail to detect some novel AOA genotypes and therefore underestimate the AOA abundances. Recently, *Nitosopumilus maritimus* SCM1, an AOA strain originating from an aquarium [58] and also present in seawater, was revealed to be able to oxidize ammonium to nitrite in the absence of oxygen [62]. Whether AOA inhabiting deeper anoxic sediment layers that are different from *N. maritimus* can do the same is still unclear. If no, anammox bacteria then may be the sole consumer of ammonium in the sediment interval between the OATZ and NATZ, the source of which is local organic matter degradation rather than diffusion from the deeper, anoxic sediments. If yes and their cell-specific rates are similar to those of anammox bacteria, the abundances of anammox in that interval are still at least one order of magnitude higher than those of AOA, making it plausible that anammox would still dominate the ammonium consumption in that interval. Therefore, in addition to the NATZ [18], anammox bacteria are likely the dominant ammonium consumers in sediments between the OATZ and NATZ.

To assess the contribution of anammox bacteria to nitrite consumption, we also quantified the contemporaneous distributions of nitrite-oxidizing and nitrite-reducing bacteria, i.e., the other two groups involved in nitrite consumption. The relative abundance of NOB affiliated with the bacterial genera *Nitrospira* and *Nitrospina* was observed to increase in the oxic zone, and then decrease to low levels (<0.5% of the total communities) in sediments without detectable oxygen (Fig. 2D). The presence of putative NOB in anoxic sediments is also supported by the calculated absolute abundances (Fig. 2H). These observations suggest that some NOB may have long longevity and can persist in anoxic sediments for long periods of time. Although NOBs are known to be metabolically versatile [e.g., as reviewed in [63]], they are not known to maintain nitrite oxidation activity in anoxic sediments. Similar to the relative abundances of *hzo* and *amoA*, the abundances of *nirS*- and *nirK*-containing nitrite-reducing bacteria are at least one order of magnitude lower than those of anammox bacteria (Fig. 2I). Despite elevated in the upper nitrite consumption zone, the abundance of *nirS* and *nirK* genes decreased to <10^3^ copies g^−1^ levels in the lower nitrite consumption zone (Fig. 2I), indicating that they may not contribute substantially to local nitrite consumption. Therefore, anammox bacteria appear to be the major players in restricting the accumulation of both ammonium and nitrite across the nitrogenous zone.

### Niche partitioning between anammox families

To elucidate the reasons leading to the two growth horizons of anammox bacteria in GC04, we examined the anammox bacteria community on the individual OTU levels (97% cutoff). Anammox bacteria were represented by 8 OTUs (OTU_2, OTU_6, OTU_180, OTU_571, OTU_595, OTU_602, OTU_4527, and OTU_4769) (Fig. 3A). Among these anammox phylotypes, only OTU_2 was detected throughout the sediment core, while the other OTUs were only detected in discrete sediment horizons (Fig. 3A). Phylogenetic analysis (Fig. 3B) indicates that OTU_2, OTU_571, OTU_602, OTU_4527, and OTU_4769 are members of the *Ca*. Scalinduaceae family, with OTU_2 matching with *Ca*. Scalindua sediminis, an anammox bacterium previously proven to be prevalent in AMOR sediments [18]. OTU_602 and OTU_4769 fell into the broad cluster containing *Ca*. S. brodae [64], *Ca*. S. profunda [22], and *Ca*. S. japonica [21], three anammox enrichment cultures from coastal sediments. The other three OTUs (OTU_6, OTU_180, and OTU_595) are members of the newly proposed anammox bacteria family *Ca*. Bathyanammoxibiaceae [23], and cluster with uncultured anammox bacteria from the AMOR area [18] and other locations such as the South China Sea [65] (Fig. 3B).

Anammox bacteria of these two families exhibit markedly contrasting distribution patterns. *Ca*. Scalinduaceae accounts for 7% of the total community in the shallowest sediment and decrease downcore until increasing again in the interval of 120–220 cm bsf, with the peak (18% of the total community) detected at 160 cm bsf (Fig. 3C). In stark contrast, *Ca*. Bathyanammoxibiaceae shows the opposite trend: it was undetectable in the shallowest most oxygenated sediment, but increases in the upper sediments to reach the peak (11% of the total community) at 50 cm bsf, before decreasing to low levels in deeper layers (Fig. 3C). Therefore, *Ca*. Bathyanammoxibiaceae is the dominant anammox bacteria family in the upper portion of the recovered core (0–75 cmbsf), while *Ca*. Scalinduaceae dominates the lower portion of the recovered core (or the extended NATZ) (110–240 cmbsf). These observations provide the first evidence of niche partitioning between the two anammox bacteria families in the marine environment.

### Population dynamics of individual anammox families and their association with the long-term niche partitioning and nitrite accumulation

Differentiating the two anammox bacteria families helps to better evaluate the population dynamics and their respective roles in nitrite consumption. The absolute abundances of these two anammox families show the same patterns as their relative abundances (Fig. 3D). The total anammox population notably does not exhibit elevated abundances around the upper nitrite consumption zone (~50 cm bsf) (Fig. 2E), which appears to indicate that the anammox bacteria population residing in this zone shrank rather than thrived in response to the elevated nitrite flux from deeper sediments. However, this decrease in the overall anammox population was caused by the fact that the decreased abundance of *Ca*. Scalinduaceae exceeded the increased abundance of *Ca*. Bathyanammoxibiaceae. The increasing abundances of *Ca*.Bathyanammoxibiaceae with depth in the upper nitrite consumption zone also suggest that anammox bacteria in this family are not decaying but growing, similar to *Ca*. Scalinduaceae in the NATZ. This evidence of net growth of anammox bacteria in the subsurface supports the results from other AMOR cores [18,23]. The peaks of the two anammox bacterial families match well with the two net nitrite consumption zones in GC04 (Fig. 3D) and also GS16-GC04 (Fig. S5B), indicating that they are likely the major contributors to the nitrite consumption in GC04: *Ca*. Bathyanammoxibiaceae is the major contributor in the upper nitrite consumption zone and *Ca*. Scalinduaceae the major one in the lower nitrite consumption zone. Additionally, the nitrite accumulation around the nitrate-depletion zone at 70 – 160 cm bsf coincides with the collapse of the combined anammox population (Fig. 2E), again suggesting the controlling role of anammox bacteria on the distribution of nitrite.

To check whether the observed anammox niche partitioning exists in other sediments, we re-analyzed the four AMOR sediment cores reported in [18]. We found similar niche partitioning in core GS16-GC04 where the porewater nitrite accumulation in the nitrate-depletion zone was also well resolved, although it seems show an opposite trend to that of GS14-GC04: *Ca*. Scalinduaceae occupies the shallower depths while *Ca*.

Bathyanammoxibiaceae dominates the deeper layers (Fig. S5). The discrepancy between these two cores may be caused by the insufficient coring depth of GS14-GC04, where only the onset of *Ca*. Bathyanammoxibiaceae in the deep NH_4_^+^-bearing sediments was captured but its potential dominance in deeper sediments was not resolved (Fig. 3D). In the other three cores reported in [18] where no nitrite accumulation was found, no clear niche partitioning between anammox bacteria of the *Ca*. Scalinduaceae and *Ca*. Bathyanammoxibiaceae can be observed. Therefore, nitrite accumulation in the AMOR sediment seems to be associated with the niche partitioning of anammox bacteria.

Low activity of microbes in subsurface sediments results in long generation times and can prolong the population evolutionary process. The observed abundance maxima of the two anammox bacteria families in GC04 were separated by ~110 cm of sedimentation (Fig. 3C and 3D). Given the sedimentation rate of ~2 cm ky^−1^ at this area [66], the duration of the niche partitioning between the two anammox family can be estimated to be about 55,000 years. The partial collapse of the whole anammox bacteria population observed during this prolonged process of niche partitioning (Fig. 2E) can be caused by the decreased supply of the two essential substrates of anammox bacteria: nitrite and ammonium. However, considering the observation that anammox bacteria abundance was higher in the low nitrite depths but lower in the high nitrite depths, it is unlikely the availability of nitrite constrained the anammox bacteria abundance. Also, the highest nitrite concentration in GC04 (3.3 μM) is close to or lower than the measured nitrite affinities of anammox bacteria [67], indicating that nitrite is unlikely inhibiting anammox bacteria in the sediment layers where nitrite accumulates. Instead, decreased ammonium supply is a plausible factor responsible for the niche partitioning. Comparing to the nitrate-depletion zone, shallower sediments may receive higher ammonium supply due to the higher organic matter degradation rates, while deeper sediments may also have higher ammonium supply due to the upward diffusion of ammonium from deeper anoxic sediments. The lower ammonium supply in the basal nitrogenous zone may have resulted in the partial collapse of the anammox population and therefore sustained the accumulation of nitrite. In other words, the two different NH_4_^+^ sources are too far apart to support anammox in the middle, which lets NO_2_^−^ accumulate.

### Potential mechanisms driving the observed anammox niche partitioning

Given the lack of anammox cultures from pelagic marine sediments, we rely on comparative genomic analysis to identify potential (and probable) reasons that lead to the niche partitioning between the two anammox bacteria families in AMOR sediments. High quality genomes are a prerequisite for such analysis. Although *Ca*. Scalindua sediminis [18] is a high-quality representative of the *Ca*. Scalinduaceae family, the previous metagenome-assembled genome (MAG) of *Ca*. Bathyanammoxibiaceae in AMOR sediments, Bin_158, was estimated to be only 74% complete [23]. Therefore, to obtain high-quality representative genomes of *Ca*.Bathyanammoxibiaceae in AMOR sediments, we performed metagenome sequencing on the sediment horizon GC05_55cm, because *Ca*. Bathyanammoxibiaceae in this sediment layer was revealed to account for 28% of the total prokaryotic community by 16S rRNA gene amplicon sequencing [23]. By metagenome assembly and binning, we obtained a high-quality MAG (96.6% completeness and 1.5% redundancy) affiliated to the *Ca*. Bathyanammoxibiaceae family. This MAG is 2.1 mega-base pairs in size, with 1905 coding genes distributed on 32 scaffolds. It has an average nucleotide identity of 98% with Bin_158 previously recovered from core GC08 [23] and therefore can be regarded as the same anammox bacterial species shown to prevail in AMOR sediments. It has a ribosomal operon, and the 16S rRNA gene (1 334 bp) is a 100% match with OTU_6 of GC04 presented here (Fig. 3B) and with OTU_23 of the four previously characterized AMOR cores [23], indicating that it can represent the most dominant Bathyanammoxibius phylotype in these AMOR cores. It also contains all necessary genes for the core anammox metabolism, including hydrazine synthase (though the alpha, beta, and gamma subunits are located at the ends of two separated contigs), hydrazine dehydrogenase, and nitrite oxidoreductase. We provisionally name this MAG *Candidatus* Bathyanammoxibius amoris (named after AMOR, the originating location of this MAG).

Using *Ca*. S. sediminis and *Ca*. B. amoris as representative genomes of the *Ca*.Scalinduaceae and *Ca*. Bathyanammoxibiaceae families, respectively, shown to dominate in this system, we performed a comparative genomic analysis to identify potential reasons that may lead to the niche partitioning between the two anammox bacteria families in AMOR sediments. The two genomes combined contain 4808 genes summarized in 1548 gene clusters, of which 917 are shared by the two genomes (Fig. 4B). Of the remaining 631 gene clusters, 457 are unique in *Ca*.S. sediminis, and the other 174 gene clusters are unique in *Ca*. B. amoris (Fig. 4B). Since both are anammox genomes, genes encoding the key enzymes of the core anammox metabolism are among the shared gene clusters (included in Supplementary dataset S2). Comparing to *Ca*. Scalindua sediminis [18], *Ca*. B. amoris lacks urease and cyanase, indicating that it does not have the capacity to conserve energy or produce extra ammonium from the degradation of urea and cyanate. Although cyanate availability in marine sediments has not been determined, urea concentrations had been measured to be 8 times lower than ammonium [68]. The majority of the urea production in anoxic sediments attributed to microbial degradation [69] of purines and pyrimidines [70], while in oxic sediments macrofauna, if exist, may also play a role in urea production [71]. The urea hydrolysis capacity may provide *Ca*. S. sediminis a competitive advantage to live in environments of limited ammonium, such as the surface oxic sediments (Fig. 3C and 3D).*Ca*.B. amoris also lacks thiosulfate reductase, an enzyme present in *Ca*. S. sediminis and also some other anammox bacteria [72] which may enable them to utilize thiosulfate as an electron acceptor. Notable unique genes present in *Ca*. B. amoris include genes encoding for lactate dehydrogenase, pyruvate:ferrodoxin oxidoreductase and [NiFe] hydrogenase, all of which may be involved in fermentation.

**Fig. 4.**
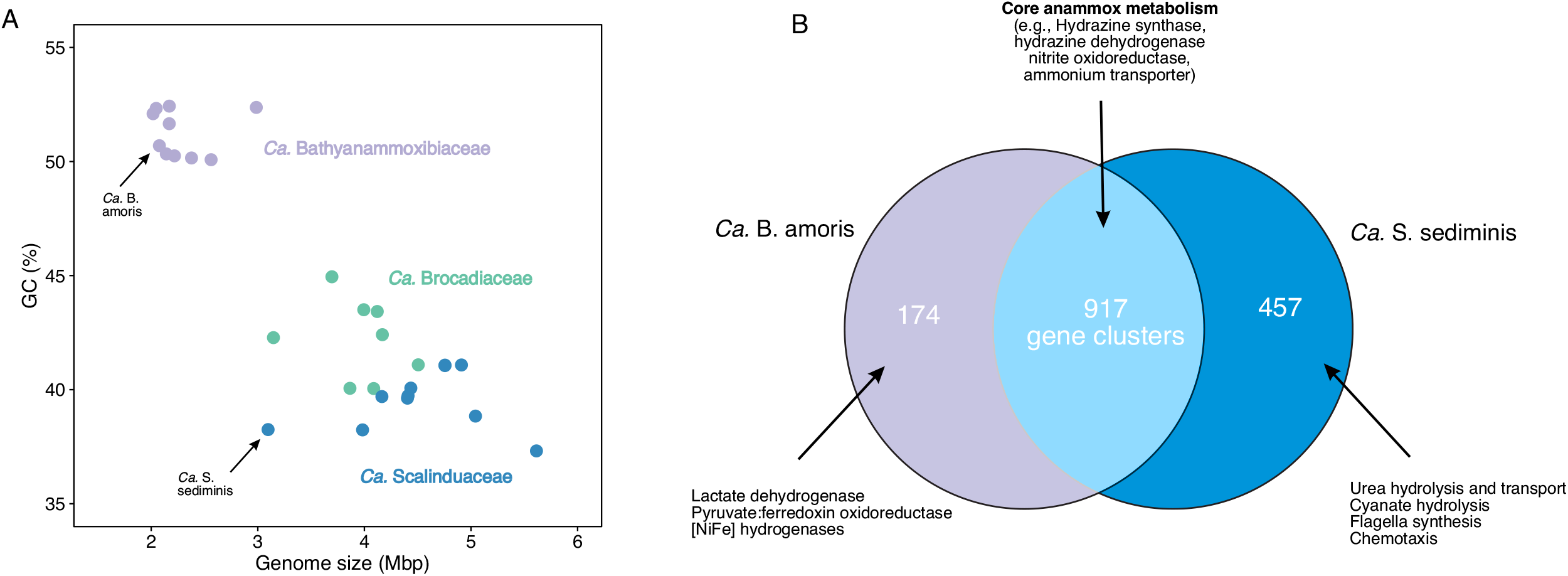
Comparative analysis of genomes of the dominant anammox bacteria in marine sediments. **(A)** A plot of genome size against GC content of the three families of anammox bacteria genomes. *Ca*. Bathyanammoxibius amoris (in this study) and *Ca*. Sediminis sediminis (Zhao et al., 2020), representatives of the families *Ca*. Bathyanammoxibiaceae and *Ca*. Scalinduaceae widespread in marine sediments, are highlighted. **(B)**Venn diagram showing the shared and unique gene clusters between *Ca*. B. amoris and *Ca*. S. sediminis.

Given that observed ammonium concentrations are profoundly different between the two niches of anammox bacteria, we investigated the types and numbers of ammonium transporters (Amt) - the essential cell apparatus for ammonium assimilation conserved in all domains of life - in available high-quality anammox bacteria genomes. We identified a total of 55 Amt among the 10 selected high-quality anammox genomes. Phylogenetic analysis of Amt suggested that anammox bacteria contain Amt of both Rh-type and MEP-type ammonium transporters (Fig. 6A). We identified one clade of anammox Amt in the Rh-type branch clustering with those of AOB and *Nitrospira* NOB [73], and 6 anammox Amt clades in the MEP-type branch (Fig. 5A). Rh-type transporter proteins in AOB [74,75] and other organisms [76] were demonstrated to have low ammonium affinity and can only be operational in high ammonium concentrations in the millimolar range, while MEP-type ammonium transporters have higher affinity [77,78] and can be efficient under conditions of low ammonium concentrations. A Rh-type Amt of low affinity is conserved in genomes of the families Brocadiaceae and Bathyanammoxibiaceae, but seem to be absent in Ca. Scalinduaceae (Fig. 5B). For the MEP-type, high-affinity Amt, anammox bacteria in the *Ca*.Scalinduaceae family have 4–8, while *Ca*. Bathyanammoxibiaceae members have only 2–5 of these ammonium transporters (Fig. 6B). Combined with the lack of access to alternative substrates and extra ammonium, encoding fewer high-affinity ammonium transporters in *Ca*.Bathyanammoxibiaceae than *Ca*. Scalinduaceae may drive the former inhabit only conditions of high concentrations or fluxes of ammonium, which is supported by the observed preference of *Ca*. Bathyanammoxibiaceae in sediment layers of (observed or inferred) higher ammonium availabilities.

**Figure 5.**
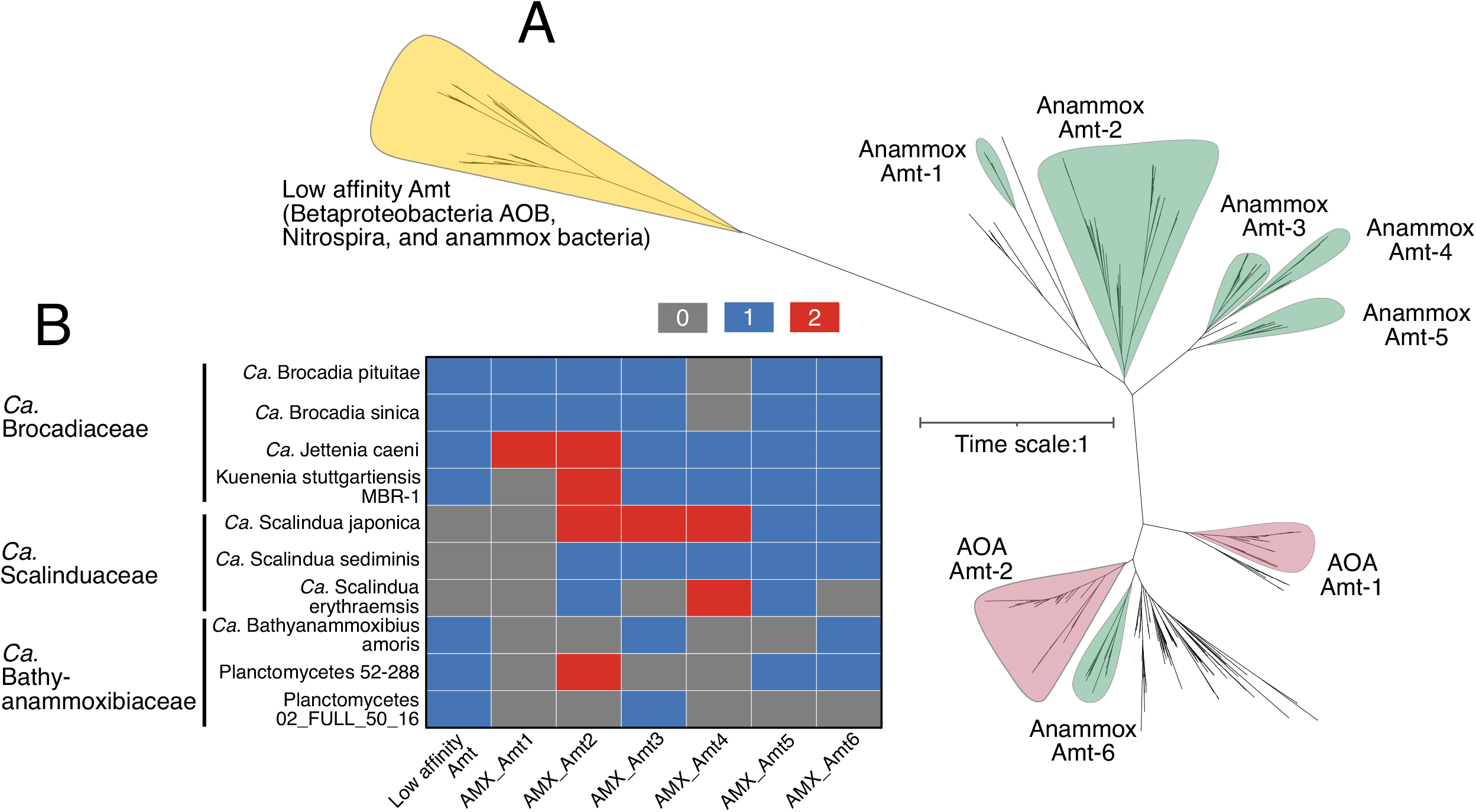
Phylogeny and distribution of ammonium transporters (Amt) in anammox bacteria. **(A)** Maximum-likelihood phylogenetic tree of Amt in anammox bacteria and other related nitrogen cycling groups (AOB, NOB, and AOA). Amt clades of nitrogen cycling groups are highlighted with different colors. The bar indicates estimated sequence divergence per residue. **(B)** Heatmap showing the occurrence of Amt in 10 selected high-quality anammox bacterial genomes.

The genome-inferred preference of higher ammonium availabilities for *Ca*. Bathyanammoxibiaceae is also consistent with the recent phylogenomic and molecular clock analysis of anammox bacteria [79]. In this work, anammox bacteria on Earth were inferred to emerge around the Great Oxidation Event [79], during ammonium is the dominant oceanic nitrogen species [1]. *Ca*. Bathyanammoxibiaceae is more deep-branching than *Ca*.Scalinduaceae, which could have more adapted to original conditions (e.g., high ammonium concentrations) of anammox bacteria.

## Conclusion

We combined biogeochemical, microbiological, and genomic approaches to study anammox bacteria and their geochemical impacts in marine sediments. We documented the prevalence of anammox in a mostly anoxic sediment core retrieved from the seafloor near the Jan Mayen hydrothermal vent fields. This core showed notable nitrite accumulation around the nitrate-depletion zone, an analogous feature also observed in 28 other globally distributed marine sediment cores and in other stratified aquatic environments. The detected anammox bacteria communities consisted of members of both families *Ca*. Scalinduaceae and *Ca*.

Bathyanammoxibiaceae. While the latter mainly thrive in the upper nitrite consumption zone, the former were detected in both the shallow oxic zone and also subsurface sediments within the NATZ. They are likely the major nitrite consumers in the zones they primarily occupy. Additionally, because of the high abundances of anammox bacteria and the low abundance of aerobic ammonia oxidizers, anammox bacteria likely also contribute substantially to the ammonium consumption in the sediment interval between OATZ and NATZ. The observed nitrite accumulation in sediment cores is accompanied by the niche partitioning between the two anammox bacteria families, which were likely driven by the differential capacities in ammonium assimilation and utilizing alternative organic nitrogen substrates like urea and cyanate. Future efforts in developing mechanistic models that can explain the observed geochemical and microbiology data while also reconciling the time history of sedimentation will greatly advance our understanding of the interactions between the nitrogen cycling processes in marine hydrothermal sediments.

## Materials and Methods

### Study area, sampling, and geochemical measurements

The core used in this study, GC04 (71°17.08’N, 6°33.69’W), was retrieved using a gravity corer from the seafloor at a water depth of 1050 meters during the CGB 2014 summer cruise onboard the Norwegian R/V G.O. Sars. This coring site is about 50 km west of the Jan Mayen hydrothermal vent field [71.2°N, 5.5°W, [24,25]] and north of the Jan Mayen fraction zone (Fig. 1A). As described in [18], the retrieved core was split into two halves on deck: one half was immediately wrapped with plastic films for archiving at 4 °C at the core repository at the University of Bergen, and the other half was used for sampling on the deck. First, the oxygen concentrations were measured using an optode by lowering the sensor into the middle part of selected depths in the working half. The optode sensors were connected to a MICROX TX3 single-channel fiber-optic oxygen meter, which was calibrated according to the manufacturer’s protocols (PreSens, Regensberg, Germany). Then, porewater was extracted using Rhizons samplers [80] from discrete depths. Microbiology subsamples were taken simultaneously with porewater extraction, using sterile 10 ml cut-off syringes from nearly identical depths as the porewater extraction, and immediately frozen at −80°C for onshore-based DNA analysis.

### Geochemical analyses

Geochemical analyses were performed using the same procedure as described in [18]. Nutrient concentrations in porewater were measured onboard. Concentrations of ammonium (NH_4_^+^), nitrate (NO_3_^−^), nitrite (NO_2_^−^), and dissolved inorganic carbon (DIC) were analyzed colorimetrically by a QuAAtro continuous flow analyzer (SEAL Analytical Ltd, Southampton, UK), following the manufacturer’s protocol. The photometric indophenol method was used for the ammonium measurement [81]. Nitrite was measured as a pink complex after reacting with *N*-1-naphthylethylenediamine dihydrochloride and sulfanilamide. The sum of porewater nitrate and nitrite was measured using the same method after reducing nitrate to to nitrite by a Cu-Cd reduction coil [82]. Nitrate concentrations were calculated as the difference between these two measurements. The protocol for DIC was based on [83]. Sulfate (SO_4_^−2^) was measured by a Metrohm ion chromatography system. Porewater samples for metal concentrations (including dissolved Mn and Fe) were acidified by ultrapure nitric acid to a final concentration of 3 vol% and stored in acid-cleaned HDPE bottles at 4°C until analysis. Metal concentrations were determined by Thermo Scientific iCap 7600 ICP-AES (inductively coupled plasma atomic emission spectrometry) at the University of Bergen. For total organic carbon (TOC) and nitrogen (TON) measurements, sediments were first dried at 95°C for 24 hours and then measured on an element analyzer (Analytikjena multi EA®4000, Jena, Germany), after inorganic carbon removal by adding 1 mL of phosphoric acid.

### Diffusive flux calculation

Diffusive fluxes of nitrate into and nitrite effluxes (both upward and downward) from the nitrate-depletion zone in sediment cores were calculated based on the measured profiles using Fick’s first law of diffusion:

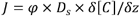

where, *J* is the flux; φ is the measured sediment porosity; *D_s_* is the sedimentary diffusion coefficient for a given solute (m^2^.yr^−1^) calculated using the *R* package *marelac* [84]; *z* is the sediment depth below the seafloor (m); and *δ[C]/δz* equals the solute (NO_3_^−^ or NO_2_^−^) concentration gradient (mmol/m^3^), calculated from nearby three data points.

### DNA extraction, PCR amplification, and sequencing

DNA for amplicon sequencing and qPCR was extracted from ~0.5 gram of sediment per sample using the PowerLyze DNA extraction kits (MOBIO Laboratories, Inc.) with the following minor modifications: 1) Lysis tubes were replaced by G2 tubes (Amplikon, Odense, Denmark), and 2) water bathed for 30 min at 60 °C before bead beating (speed 6.0 for 45 seconds) using a FastPrep-24 instrument (MP Biomedicals). A blank extraction (without sediment addition) was carried out in parallel with the sample extraction batch following the same procedure. The DNA was eluted into 80 μL of molecular grade double-distilled H_2_O (ddH_2_O) and stored at −20 °C until analysis. 16S rRNA genes were amplified using the primer pair 515F/806R in a two-round amplicon preparation [18], with an optimal PCR cycle number in the first round for each sample to minimize over-amplification. Amplicon libraries were sequenced on an Ion Torrent Personal Genome Machine.

### Amplicon sequencing data analysis

Sequencing reads were quality filtered and trimmed to 220 bp using the USEARCH pipeline [85] and chimera were detected and removed using UCHIME. Trimmed reads were clustered into operational taxonomy units (OTUs) at >97% nucleotide sequence identity using UPARSE [86]. Most of the OTUs detected in the extraction blanks (negative controls) were manually removed, except for a few OTUs that may be introduced into the blanks by cross-contamination. Overall, >99.9% of reads in the negative controls were removed. Samples were subsampled to 20,000 reads for each sediment horizon. The taxonomic classification of OTUs was performed using the lowest common ancestor algorithm implemented in the CREST package [87] with the SilvaMod128 database as the reference. The relative abundance of anammox bacteria was taken as the total percentage of the OTUs affiliated with the families *Ca*. Scalinduaceae and *Ca*. Bathyanammoxibiaceae [23]. The distribution of individual anammox OTUs was visualized in heatmaps generated using the *R* package *ggplot2* [88].

### Quantification of 16S rRNA genes and functional genes

Abundances of anammox bacteria were quantified using qPCR by targeting the *hzo* gene (encoding the hydrazine dehydrogenase responsible for the degradation of hydrazine to N_2_) using the primer pair hzoF1/hzoR1 [89] following the procedure described elsewhere [35]. The abundances of denitrifying bacteria were quantified by targeting the *narG* (coding the periplasmic nitrate reductase alpha subunit), *nirS* and *nirK* genes (coding cytochrome *cd1*-and Cu-containing nitrite reductases, respectively), using the protocol described in [35]. In addition, archaeal and bacterial 16S rRNA genes were quantified as described in [90]. Total cell abundance was estimated from 16S rRNA gene copies, assuming a single copy of 16S rRNA genes in each bacterial or archaeal genome [26]. All gene abundances were determined triplicate in qPCR, and standard deviations are presented using horizontal error bars. Absolute abundances of the aforementioned groups were also calculated as the product of the total cell abundance and the percentage of these groups in the total community assessed by amplicon sequencing.

### Metagenome sequencing, assembly, and binning

To recover high-quality genomes of *Ca*. Bathyanammoxibiaceae, we focused on the sediment horizon of 55 cm of core GS16-GC05, because our previous survey indicated that this particular sediment horizon harbors the highest relative abundance of *Ca*. Bathyanammoxibiaceae in the total archaea and bacteria community [23]. We extract the total DNA from 6.4 g of sediment (~0.4-0.6 g sediment in each of the 12 individual lysis) using PowerLyze DNA extraction kits (MOBIO Laboratories, Inc.) following the manufacturer’s instructions. The DNA extracts were iteratively eluted from the 12 spin columns into 100 μL of ddH_2_O for further analysis.

Shotgun metagenome libraries were constructed using a NEBNext Ultra II FS DNA Library Prep Kit (New England Laboratories) and sequenced (2×150 bp paired-end) on an Illumina NextSeq 500 sequencer at the MIT BioMicro Center. The quality of the reads and the presence of adaptor sequences were first checked using FastQC v.0.11.9 [91]. Adapters were removed and reads trimmed using BBDuk implemented in the BBMap package (Bushnell, 2014). The overall quality of processed reads was evaluated in a final check with FastQC v.0.11.9 [91], to ensure only high-quality reads were used in the downstream analysis. The quality-controlled reads were *de novo* assembled into contigs using Megahit v.1.1.2 [92] with the k-mer length varying from 27 to 117 and a contig length threshold of 1000 bp. Contigs were grouped into genome bins using three programs: MaxBin2 v2.2.6 [93], MetaBAT v2.15.3 [94], and CONCOCT v1.1.0 [95], all with the default parameters. The resulting bins from these three programs were subject to dereplication and aggregation by DASTool v1.1.4 [96] with the default parameters. The quality of the obtained genome bins was assessed using the option “lineage_wf” of CheckM v.1.1.3 [97]. To improve the quality of the genomes affiliated to the Brocadiales order, the quality-controlled reads were mapped onto the contigs using BBmap (Bushnell, 2014), and the mapped reads were re-assembled using SPAdes v.3.14.0 [98]. After removal of contigs shorter than 1000 bp, the resulting scaffolds were visualized and re-binned manually using gbtools [99] as described in [18]. The quality of the resulting *Ca*. Bathyanammoxibius genome was checked using the CheckM “lineage_wf” command again, based on the Planctomycetes marker gene set.

### Comparative genomic analysis

We performed a comparative analysis on the genomes *Ca*. Scalindua sediminis [18] and *Ca*.Bathyanammoxibius amoris (recovered in this study), the dominant species of the two anammox bacteria families in marine sediments [23]. We did the analysis using Anvio v.7.1 [100] according to the workflow described at http://merenlab.org/2016/11/08/pangenomics-v2/. All genomes were first annotated using Prokka v.1.14 [101] and BLASTp using the Clusters of Orthologous Groups of proteins (COG) [102] as the reference database. The comparative genomic analysis use BLAST to quantify the similarity between each pair of genes, and the Markov Cluster algorithm (MCL) [103] (with inflation parameter of 2) to resolve clusters of homologous genes. The shared and unique genes in the two genomes were identified via the functional enrichment analysis [104].

### Phylogenetic analyses

A maximum-likelihood phylogenetic tree based on 16S rRNA genes was reconstructed for known anammox bacteria and close relatives of the putative anammox OTUs identified via BLASTn [105] in the NCBI database. Sequences were aligned using MAFFT-LINSi [106] and the maximum-likelihood phylogenetic tree was inferred using IQ-TREE v. 1.5.5 [107] with GTR+F+R5 as the best-fit substitution model selected by ModelFinder [108]. 1000 ultrafast bootstraps iterations were performed using UFBoot2 [109] to assess the robustness of tree topology.

For the phylogeny of Amt (ammonium transporter), the sequence of anammox genomes were extracted from the Prokka annotation and used as the queried in BLASTp [105] searches against the NCBI database (>50% similarity were retained), to identify its close relatives. These sequences were complemented with known nitrifiers (e.g. ammonia-oxidizing bacteria (AOB) from the genera of Nitrosospira, Nitrosomonas, Nitrososcoccus, nitrite-oxidizing bacteria (NOB) from Nitrospira and Nitrospina, and ammonia-oxidizing archaea (AOA) from the Thaumarchaeota phylum) and aligned using MAFF-LINSi [106]. The alignment was trimmed using trimAl [110] with the mode of “automated”. A maximum likelihood phylogenetic tree was reconstructed using IQ-TREE v. 1.5.5 [107] with the LG+F+R7 as the best-fit substitution model and 1,000 ultrafast bootstraps.

## Data availability

All sequencing data used in this study are available in the NCBI Short Reads Archive under the project number PRJNA854201. Raw metagenome sequencing data of GC05_55 cm is available in the NCBI database under the BioSample number SUB11625283. The genome sequence of *Ca*. Bathyanammoxibius amoris is available under the accession number JAMXCW000000000. Raw geochemical data can be found in the SI Appendix, Dataset S1.

## Acknowledgments

Sediment coring opportunities from the AMOR area were made possible by the chief scientist Rolf Birger Pedersen and the crew of R/V G.O. Sars. We thank Anita-Elin Fedøy for the amplicon preparation, Michael Melcher and Steffen Lydvo for sampling collection and DNA extraction, and Jan-Kristoffer Landro for sediment carbon and nitrogen contents measurements. This work was funded by the Research Council of Norway through the Centre for Excellence in Geobiology, the K.G. Jebsen Foundation and Trond Mohns Science Foundation to S.L.J., and Simons Foundation grant 622065 and National Science Foundation grants OCE-2138890 and OCE-2142998 to A.R.B. R.Z. is supported by the MIT Molina Postdoctoral Fellowship.

## Author contribution

R.Z. and S.L.J. conceived the study. R.Z., D.R., I.H.T, and S.L.J. collected samples onboard the cruise. D.R. and I.H.T. performed the porewater extraction and analysis. R.Z., S.L.J, and A.R.B. collected and analyzed the genomic data. R.Z. performed the DNA analyses and interpreted the results. R.Z. and A.R.B. wrote, and all authors edited and approved the manuscript.

