## Supplementary Information for "Nitrite accumulation and the associated anammox bacteria niche partitioning in marine sediments"

**Supplementary Text**

***Note 1: The most surface sediment may have been lost during coring***

It is also worth noting that (i) a low concentration of oxygen (15 µM), (ii) a high concentration of nitrate (31.3 µM), and (iii) a high concentration of DIC (2.75 mM) were measured in the most surface sediments, the combination of which suggested that the most top sediments were very likely lost during coring. The exact length of the lost core is difficult to determine based on reconciling geochemical profiles from multiple cores, because so far there is only one core was recovered from this area. However, using the same gravity coring devise, previous studies have reported < 17 cm of core-top lost from deep-sea sediments (Chong et al., 2018; Lund et al., 2018).

**Supplementary Figures**

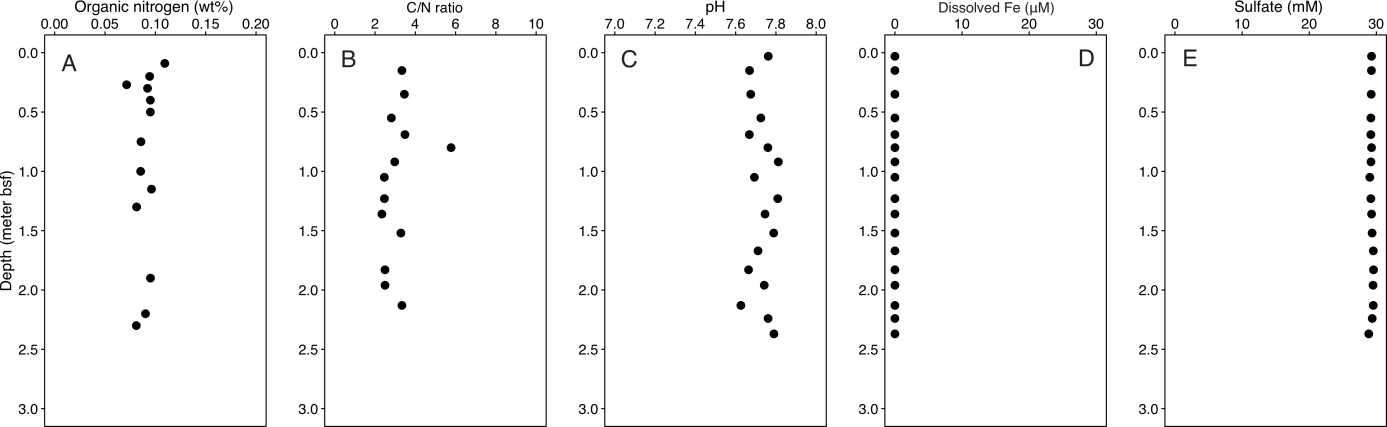

**Fig. S1. Organic nitrogen content (A), organic C/N ratio (B), porewater pH (C), dissolved Fe concentration (D), and Sulfate concentration (E) measured in GC04.**

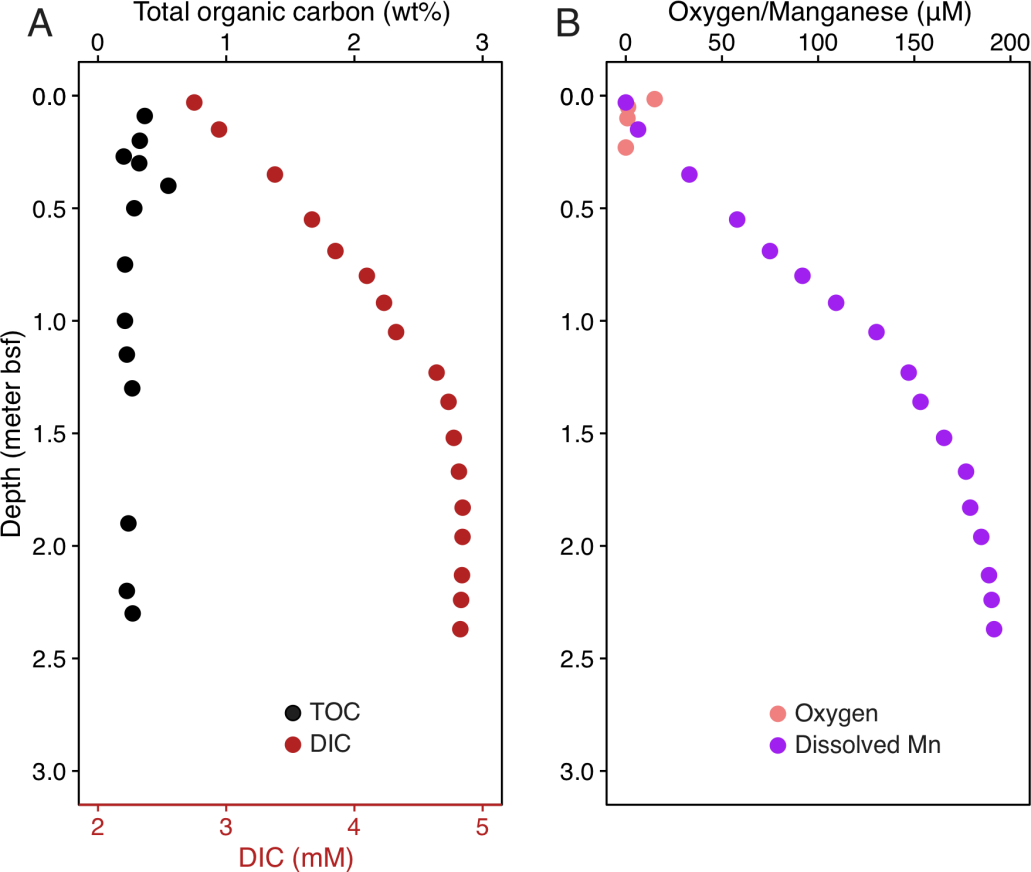

**Fig. S2. Geochemical profiles of GC04.** Depth profiles of total organic carbon (TOC) and dissolved inorganic carbon (DIC) (**A**), oxygen and dissolved Mn (**B**).

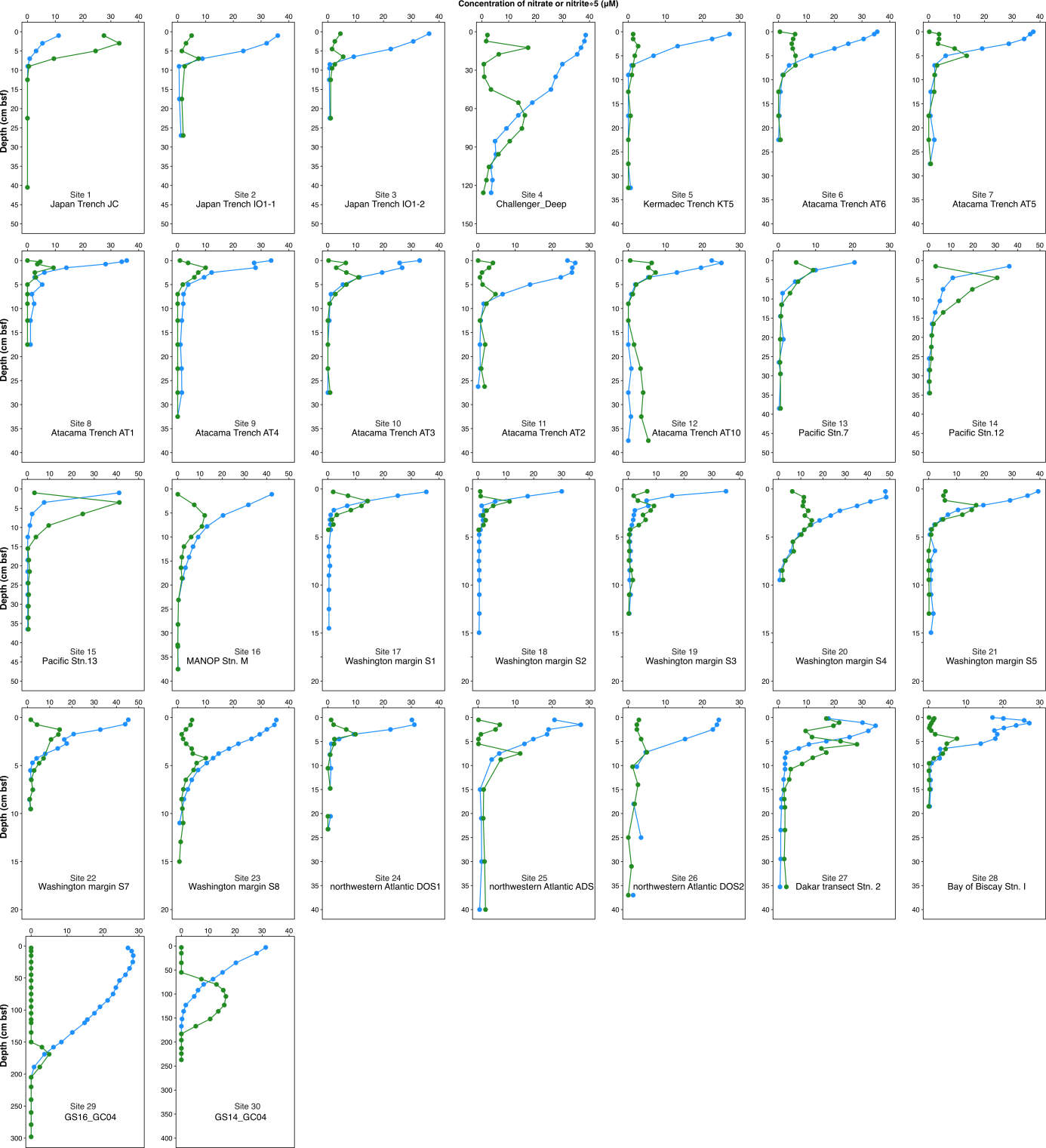

**Fig. S3. Porewater profiles of nitrate (blue) and nitrite (×5, green) in the porewater of marine sediment cores where nitrite accumulation in the nitrate-depletion zone was apparent.**

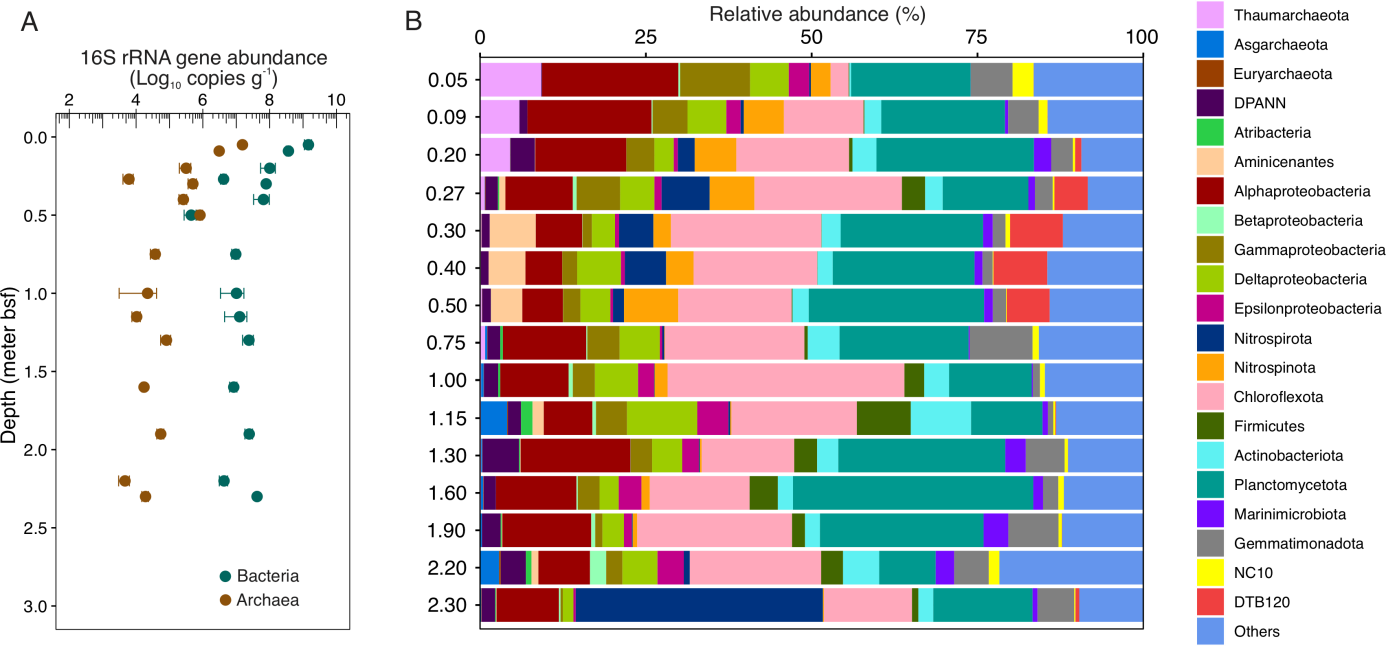

**Fig. S4. Abundance (A) and community structure (B) of archaea and bacteria in GC04. (A)** Abundances of archaea and bacteria determined by qPCR using domain-specific primers**.** Dots show the mean values and error bars denote the standard deviations of triplicate measurements. **(B)** Community structure assessed by 16S rRNA gene amplicon sequencing. Minor taxa (with relative abundances of <1%) were grouped into the category “Others”.

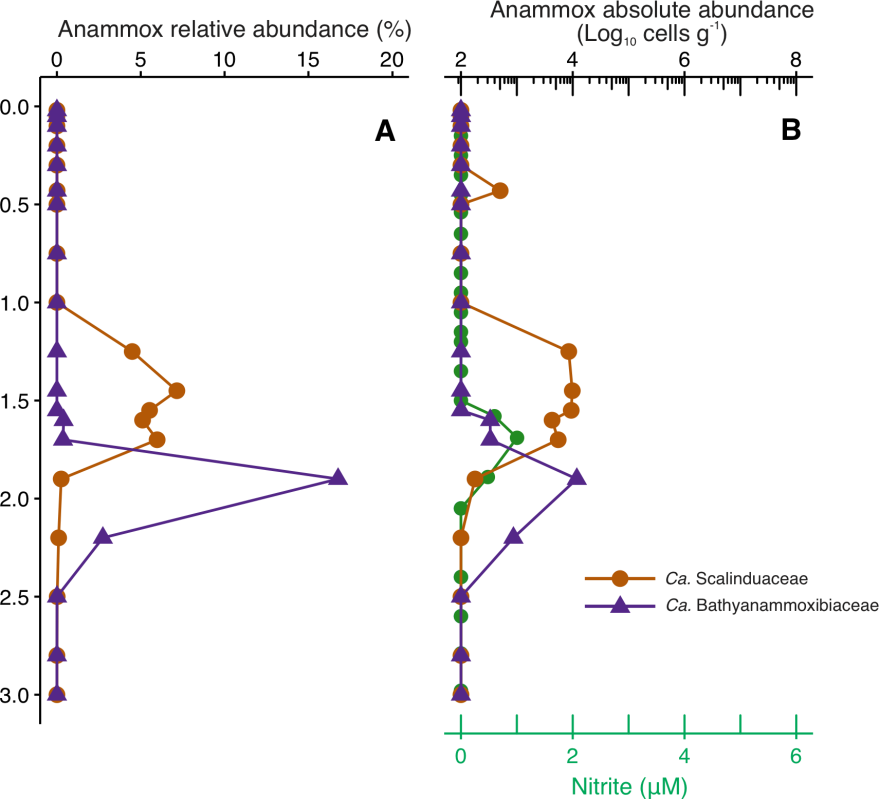

**Figure S5. The distribution of anammox bacteria families and porewater nitrite in cores GS16-GC04 (A, B) and GS16-GC05 (C, D).** The nitrite profile in (B) is the same as that shown in Fig. 2C.
